# Phenomics data processing: Extracting dose-response curve parameters from high-resolution temperature courses and repeated field-based wheat height measurements

**DOI:** 10.1101/2021.07.23.453040

**Authors:** Lukas Roth, Hans-Peter Piepho, Andreas Hund

## Abstract

Temperature is a main driver of plant growth and development. New phenotyping tools enable quantifying the temperature response of hundreds of genotypes. Yet, for field-derived data, temperature response modeling bears flaws and pitfalls concerning the interpretation of derived parameters. In this study, climate data from five growing seasons with differing temperature distributions served as starting point for a growth simulation of wheat stem elongation, based on a four-parametric temperature response function (Wang-Engel) including all cardinal temperatures. In a novel approach, we re-extracted dose-responses from the simulation by combining high-resolution (hours) temperature courses with low-resolution (days) height data. The collection of such data is common in field phenotyping platforms. To take advantage of the lack of supra-optimal temperatures during the stem elongation, simpler (linear and asymptotic) models to predict temperature-response parameters were investigated. The asymptotic model extracted the base temperature of growth and the maximum absolute growth rate with high precision, whereas simpler, linear models failed to do so. Additionally, the asymptotic model provided a proxy estimate for the optimum temperature. However, when including seasonally changing cardinal temperatures, the prediction accuracy of the asymptotic model was strongly reduced. In a field study with three winter wheat varieties, significant differences were found for all three asymptotic dose-response curve parameters. We conclude that the asymptotic model based on high-resolution temperature courses is suitable to extract meaningful parameters from field-based data.

## 1. Introduction

As early as 1984, it was noted that the validity and scope of a crop growth model depends on the available data, which are, however, severely limited (Porter and Rayner, 1984). Today, high-throughout field phenotyping (HTFP) potentially enables to quantify crop growth—and consequently crop response—of hundreds of genotypes under field conditions (Araus et al., 2018; Tardieu et al., 2020). Understanding crop responses to environmental factors is essential to secure global food demands: Improvement in wheat yields stagnated on a global scale in the past three decades (Brisson et al., 2010; Laidig et al., 2017). It was suspected that for Europe, the changing climate increasingly impacts wheat yields (Brisson et al., 2010). Mitigating these changes in plant breeding to improve yield requires understanding crop responses to environmental factors (Ramirez-Villegas et al., 2015). A main driver of plant growth and development is temperature (Porter and Gawith, 1999).

### 1.1. Approaches to quantify temperature responses under controlled conditions

The effects of changes in temperature on crops are well studied in controlled environments, but the translation of insights to the field is not straightforward. The temperature response of developmental processes is usually studied by exposing plants to different temperature regimes—often during rather short time period of their life cycle. Two different approaches are used: one can either (1) expose different plants of the same genotype to different temperatures (Slafer and Rawson, 1995; Hund et al., 2012; Reimer et al., 2013), or (2) apply short phases of different temperatures to the same plant. Examples of the latter approach are the studies on leaf growth performed in the indoor platform ‘Phenoarch’, summarized by Parent and Tardieu (2012). This approach requires that the temperature response is measured at a constant growth phase, during which growth is linear when the temperature remains constant. Hence, the response-pattern can be observed non-destructively on the same plant, offering a high throughput. In such experiments, the conditions of all but one covariate may be kept constant. Furthermore, the reversibility of the process can be tested by repeatedly applying the same conditions during the course of the experiment.

In general, developmental processes follow some sort of peak function in response to increasing temperatures, i.e., there is a base temperature *T*_min_ at which growth starts, an optimal temperature *T*_opt_ at which growth reaches its maximum absolute growth rate *r*_max_, and a maximum temperature *T*_max_ at which growth stops (Porter and Gawith, 1999). Depending on the research field, different models are used to approximate such functions— for an overview see, e.g., Wang et al. (2017); Parent et al. (2019) for plants, and Rebaudo et al. (2018); Rebaudo and Rabhi (2018) for ectoterms (i.e., animals whose regulation of body temperature depends on external sources). In their work, Wang et al. (2017) distinguish four types of temperature response functions with increasing complexity: linear functions with no optimum or maximum temperature (Type 1), capped linear or curvilinear functions with an optimum but no maximum temperature (Type 2), segmented linear functions with optimum and maximum temperatures (Type 3), and segmented linear functions or curvilinear functions with three cardinal temperatures (Type 4). Clearly, functions of Type 4 are most appropriate to describe temperature responses. Among the most widely used Type 4 functions for plants are the Wang-Engel cardinal temperature function (Wang et al., 2017) and a modified version of the reaction rates equation from Johnson et al. (1942) applied to different growth processes in a variety of major crops by Parent and Tardieu (2012). We will refer to these different response functions in detail further down.

### 1.2. Cardinal temperatures of wheat

In their meta-analyses reporting cardinal temperatures of different developmental processes of wheat, Porter and Gawith (1999) summarized 65 controlled condition and field studies with regard to the minimum (*T*_min_), optimum (*T*_opt_) and maximum (*T*_max_) temperature. Mean cardinal temperatures (based on daily averages) for shoot growth were 3.0 °C (*T*_min_), 20.3 °C (*T*_opt_) and > 20.9 °C (*T*_max_). Notably, *T*_opt_ and *T*_max_ were very close together in many studies.

While Porter and Gawith (1999) were reporting on cardinal temperatures related to the phenology of wheat, Parent and Tardieu (2012) were relating to those of a single organ, specifically, a wheat leaf. In contrast to the comparably low *T*_opt_ summarized by Porter and Gawith (1999), Parent and Tardieu (2012) reported a significantly higher *T*_opt_ of 27.7 °C when summarizing eight studies and own data. Since a wheat stem is composed of several growing internodes, we consider its (long-term) elongation to be closer related to phenology than to (short-term) organ growth. Consequently, we will use the more conservative, i.e., lower, temperature optimum reported by Porter and Gawith (1999) in this study.

Yet, in addition to the short-term response of growing organs within the borders of genotype-specific cardinal temperatures, there is a long-term adjustment of these cardinal temperatures during the growing season. Slafer and Rawson (1995) examined *T*_min_ and *T*_opt_ for leaf appearance rates of vernalized winter wheat plants within three phases, i) after vernalization up to the terminal spikelet, ii) from the terminal spikelet up to heading, and iii) from heading to anthesis. The authors found that, averaged across cultivars, *T*_min_ was −1.9 °C, 1.2 °C and 8.1 °C, while *T*_opt_ was < 22 °C, 25 °C and > 25 °C for the three different phases, respectively. In addition, the cultivars differed substantially for the two parameters. Consequently, one may assume that in particular the base temperature may change throughout the season in a cultivar-specific manner. Nevertheless, the estimates from Slafer and Rawson (1995) of cardinal temperatures and their changes were derived from a controlled environment study. The benefits of controlled environment phenotyping lies in the ability to test a wide range of temperature regimes while keeping other influential factors, such as vapor pressure deficit and light, constant. However, plants usually do not experience such conditions in the field, as discussed by Slafer and Rawson (1995).

### 1.3. Approaches to quantify temperature responses with high-throughput in the field

Under field conditions, the temperature regimes are distinctly different from the controlled-environment studies considered so far. The environmental covariates in the field follow an annual and diurnal pattern to which plants adjust their life cycle. HTFP offers the possibility to assess crop growth under such more realistic conditions with high temporal and spatial resolution (Rebetzke et al., 2019). Hence, if assuming that temperatures measured at a local weather station are closely related with effective in-canopy temperatures—i.e., meristem temperatures (Gallagher et al., 1979)—one can use the temperature course during the season to quantify the genotype-specific temperature responses. Still, such studies are sparse. Grieder et al. (2015) extracted the base temperature and response to increasing temperatures at early growth stages before terminal spikelet formation. The authors were able to extract heritable slopes and intercepts of linear responses. Kronenberg et al. (2020a) were able to extract heritable response traits for the stem elongation phase based on data from three seasons. Again, the authors used linear regressions of growth versus temperature as model. Kronenberg et al. (2020a) argued that the recorded average temperatures in the measuring intervals were well below the optimum temperature of 27.7 °C reported by Parent and Tardieu (2012), which would justify approximating a curvilinear with a linear function. However, the optimum temperature of 20.3 °C summarized by Porter and Gawith (1999) is distinctly lower. Following those results, it is likely that growth reached its optimum at several hours of the day during their measurement period. Therefore, the extracted temperature response (slope) may be greatly affected by the optimum temperature and maximum abolute growth rate at this temperature. Thus, modelling plant response to temperature based on HTFP-derived data poses several challenges, e.g., the comparably sparse measuring frequency, the unpredictable range of temperatures, and the changes of cardinal temperatures during the measuring period.

### 1.4. Research aims

HTFP data are characterized by irregular and long trait measurement intervals but regular and short covariate measurement intervals from a weather station. Our main research question was how to parameterize genotype-specific temperature response pattern from such HTFP data. Using the stem elongation phase of winter wheat as an example, we aimed to evaluate a new modelling approach with regard to its suitability. Therefore, we simulated the temperature-response in the stem elongation phase using a Wang-Engel response function. The chosen parameters were based on existing literature on cardinal temperatures in winter wheat (Porter and Gawith, 1999) and enhanced with realistic temperature recordings of the stem elongation phase from a local weather station at the field phenotyping platform site of ETH Zurich (FIP) (Kronenberg et al., 2020a). To evaluate the strengths and limitations of our approach, we added changing cardinal temperatures, shifts in the start and end of the elongation phase, measurement noise, and spatial field inhomogeneities to our simulation (Appendix, Figure 6).

## 2. Background

Modeling the stem elongation phase of winter wheat on a HTFP trait level (i.e., plant heights) requires assuming a theoretical whole-plant mechanism that is the result of the combination of a vast number of underlying physiological mechanisms (Tardieu et al., 2020). Although the number of combinations of such mechanism may be near-infinite, Tardieu et al. (2020) argued that natural selection and breeding activities have constrained them to a limited number, resulting in a ‘meta-mechanism’ that is consistent across modeling scales. In the following, we provide background information on tools that allow extracting parameters of the ‘meta-mechanism’ temperature response from HTFP data.

### 2.1. Parametric versus semi-parametric approaches

Semi-parametric approaches (e.g., using smoothing techniques) are often favored to extract traits related to growth processes from HTFP time series (van Eeuwijk et al., 2019; Roth et al., 2021a; Pérez-Valencia et al., 2022), as they avoid the daunting task of modeling explicitly the—potentially unknown—influence of time-dependent covariate courses. In a previous research, we have shown that semi-parametric approaches are suitable to extract timing of key stages and quantities at defined time points or periods from simulated HTFP data. Those data were generated using a dose-response curve that was unknown to the model being fitted, and corresponding covariate dependencies masked by a “smoothing” part in the model (Roth et al., 2021b).

In contrast, to extract temperature responses, one cannot avoid modeling a dose-response curve explicitly, as one is interested in its parameters. Consequently, we hereafter propose a parametric approach. Although our focus is on HTFP data, the approach is also directly applicable to indoor platform data, but additionally considers field-specific characteristics.

In comparison to greenhouse or climate chamber data, HTFP data have two major drawbacks: (i) Field-based measurements are notoriously noisy due to environmental and soil inhomogeneities and measurement errors; (ii) the suitability of measured trait time series and covariate courses to extract dose-response curves requires large numbers of years and/or locations or the good luck of the scientist to catch at least one year with a (yet unknown) close-to-optimal distribution of temperatures along the season. Research in agriculture has already addressed the issue of spatial heterogeneity with the development of highly specialized experimental designs (e.g., by augmenting row-column designs with check cultivars (Piepho and Williams, 2016)) and matching statistical analysis (e.g., by adding spatial components to the model (Piepho and Williams, 2010)). We could already show that following these principles is of advantage for HTFP as well (Roth et al., 2021b). For brevity, we therefore focus on the extraction of response traits from time series but refrain from further processing to adjusted genotype means. To allow determining and comparing the performance of non-linear and linear models and the requirements on an ‘close-to-optimal’ distribution of temperatures along the season, we use a simulation that is parametrized based on real field data.

### 2.2. Temperature response functions

The linear temperature response model used in Grieder et al. (2015) is defined as

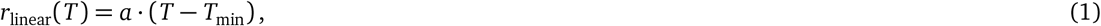

where *T* is the temperature, *a* is the slope of the response, and *T*_min_ the base temperature. Note that such a model causes negative growth rates *r* for *T* < *T*_min_ and does not include an optimum temperature *T*_opt_. Therefore, it is likely that the extracted temperature response (slope) is affected by the optimum temperature and maximum absolute growth rate at this temperature. Additionally, using such a Type 1 response (a linear function with no optimum or maximum temperature, Wang et al. (2017)) will come to its limitation when data ranges span a whole growing season where temperatures also extend to supra-optimal ranges (Weikai and Hunt, 1999; Parent et al., 2019). In the crop model community, models describing the response to temperature vary widely, but most of them consider an optimum temperature beyond which the growth rate levels off (Bonhomme, 2000; Parent et al., 2010; Wang et al., 2017).

Arrhenius type functions were shown to adequately describe the dose-response relationship between temperature and growth across a wide range of temperatures and species (Parent et al., 2010; Parent and Tardieu, 2012). However, it was disputed whether this approach allows for a mechanistic interpretation of the different model parameters (Clavijo Michelangeli et al., 2016). Evidently, research in wheat so far mainly focused on determining (non-mechanistic) cardinal temperatures such as the lower and upper temperature of growth *T*_min_ and *T*_max_ and the optimum temperature *T*_opt_ where the growth rate *r* is maximal (Porter and Gawith, 1999). Wang et al. (2017) could show that using the Wang-Engel temperature function (Wang and Engel, 1998) allows to adequately replace an Arrhenius type functions to describe growth rates using these cardinal temperatures,

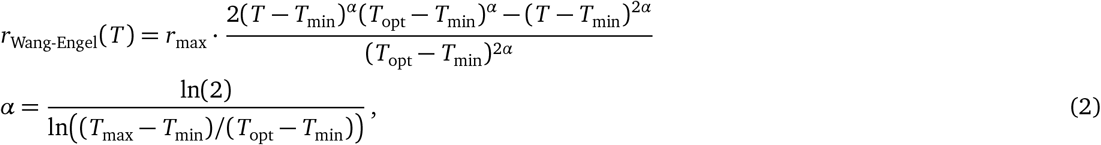

where *T* is the temperature, *r*_max_ the maximum absolute growth rate at the temperature optimum *T*_opt_, *T*_min_ the lower base temperature and *T*_max_ the upper base temperature of growth.

The Wang-Engel temperature function is based on three cardinal temperatures *T*_min_, *T*_max_ and *T*_opt_. If aiming to extract genotype-specific cardinal temperatures as well as the maximum absolute growth rate *r*_max_ from HTFP data, this would require to fit a four-parametric non-linear model with parameters that have interdependent constrains (e.g., *T*_min_ < *T*_opt_). In our experience, extracting realistic parameter values by fitting such a model to noisy HTFP data is practically impossible for different reasons, i.e., (i) the optimum temperature *T*_opt_ and maximum absolute growth rate *r*_max_ are closely correlated, (ii) for field experiments performed in target environments (without stress treatments) in temperate climates, the maximum temperature *T*_max_ is typically outside the range of observed data, and (iii) proximity of cardinal temperatures *T*_min_ and *T*_opt_ may make it challenging to achieve convergence. Some authors have reacted to these restrictions by fixing one or multiple parameters and normalizing growth rates (Parent and Tardieu, 2012).

As a previously unconsidered alternative, we propose a Type 2 response that integrates an optimum temperature (Wang et al., 2017) to approximate the Arrhenius type functions of Parent and Tardieu (2012) based on an asymptotic model (Figure 1c),

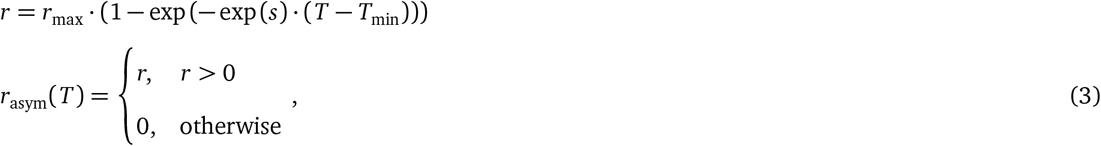

where *T* is the temperature, *r*_max_ is the maximum absolute growth rate (and therefore the asymptote of the curve), *T*_min_ the base temperature where the growth rate is zero, and *s* characterizes the steepness of the response (natural logarithm of the rate constant, thus ‘*lrc*’) (Pinheiro and Bates, 2000). Please note that (1) *lrc* is a parameter in our model and not—as the name may indicate—a constant, (2) Equation 3 reduces to a monomolecular / Mitscherlich growth function if replacing the first factor *r*_max_ with one and substituting *s* with ln(*r*_max_) (Heinen, 1999). The proposed asymptotic model does not consider a maximum temperature and therefore supra-optimal range, and has no inflection point (Paine et al., 2012). It requires fitting a three-parametric non-linear model. Monomolecular growth functions such as the asymptotic function were frequently used to describe growth rates in proportion to a substrate level, e.g., in response to photon flux density (Causton and Dale, 1990). Nevertheless, we are not aware of previous applications to temperature response problems in plant science. We would like to note that such an approach is only appropriate if one expects the maximum temperature *T*_max_ to be outside the range of observed data, as it is the case for the stem elongation phase of winter wheat—at least under the temperate climate of Switzerland.

**Figure 1:**
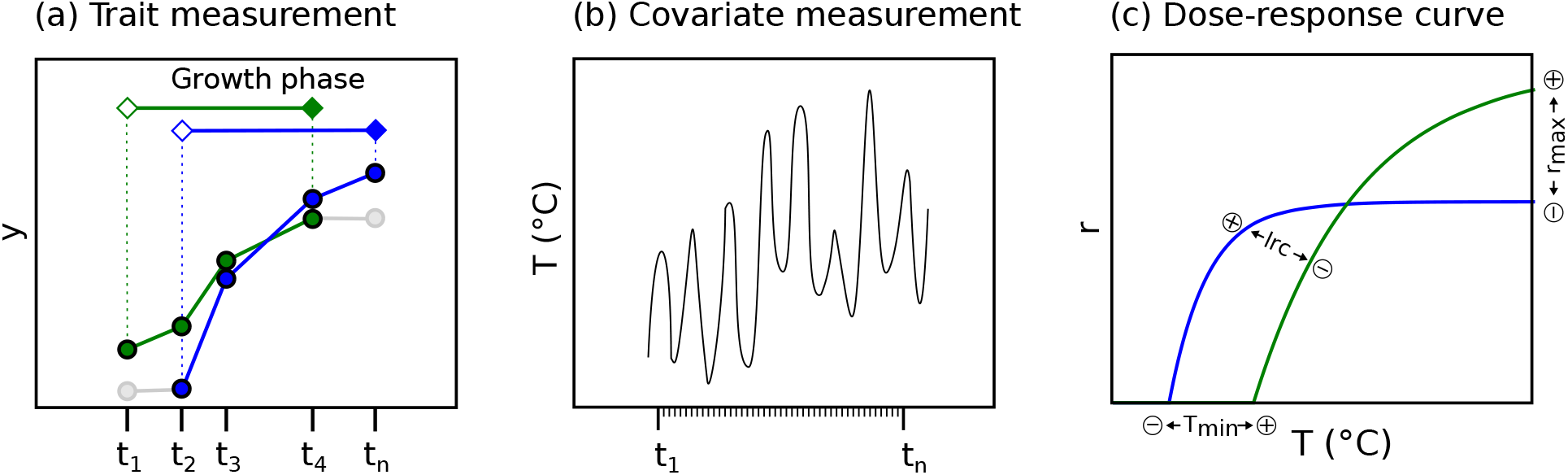
Schematic drawing of trait measurements with irregular and long intervals for two contrasting genotypes (green and blue) in their growth phase (solid) and outside their growth phase (grey) (a), covariate measurements with short and constant intervals (b), and the asymptotic dose-response model based on a maximum absolute growth rate (*r*_max_), minimum temperature (*T*_min_), and steepness of the response (*lrc*) (c).

### 2.3. Combining irregular and long trait measurement intervals (days) with constant and short covariate measurement intervals (hours)

Independent of the specific dose-response model, one can describe a phenotype *y* at time point *t* (Figure 1a) as the result of such a dose-response model *r*() (Figure 1c) dependent on a temperature course *T_t_* (Figure 1b) and a genotype-specific set of crop model parameters *θ*, e.g., *θ* = (*T*_min_, *T*_opt_, *r*_max_),

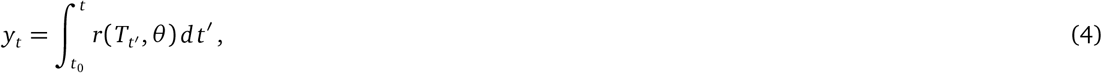

where *t*_0_ is the time point were growth started and therefore a timing of key stage trait that needs to be determined beforehand. Trait and covariates such as canopy height and temperatures are only measured at certain points in time, dependent on the measurement interval Δ*m*. We therefore may discretize Equation 4 to

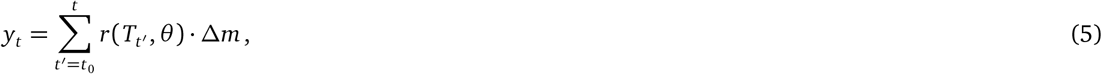

where *t′* are measurement time points with (*t′* = *t*_0_, *t*_0_ + Δ*m*, *t*_0_ + 2Δ*m*,…, *t*), e.g., days.

Note that fitting Equation 5 to data implies the same measurement interval for traits *y_t_* and temperatures *T_t′_*. Although the goal of HTFP is to measure at small measurement intervals, unfavorable weather conditions may prevent regular measurements at all times, while registering covariates (e.g., temperature) is weather-independent and done with constant high frequency. Outdoor phenotyping therefore produces trait measurements with irregular and long measurement intervals Δ*m_d_* (e.g., measurements every 3–4 days) but covariate measurements with constant and short measurement intervals Δ*m_h_* (e.g., hourly measurements). Therefore, before applying Equation 5 to data, an aggregation step is required.

A common technique is to align trait and covariate measurements by aggregating the covariate values to a mean (e.g., Kronenberg et al., 2017, 2020a),

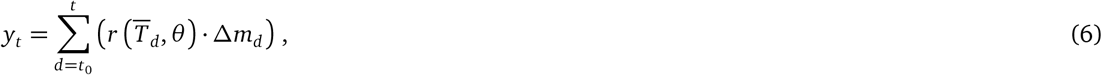

where 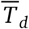 is the mean of all covariate measurements made in the time period between successive trait measurement time points *d* where (*d* = *t*_0_, *t*_0_ + Δ*m*_1_, *t*_0_ + Δ*m*_1_ + Δ*m*_2_,…, *t*). In this study, we propose an alternative approach: One may also apply the dose-response function *r* to each covariate value *T_dh_* where *h* indexes covariate values between successive trait measurement time points *d*, (*h* = 1,…, *n_d_*), and sum up the responses to the trait measurements,

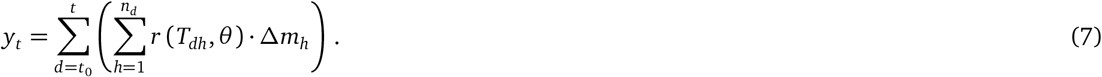

In the following, the first method is denoted the *T*_mean_ method and the second method the *T*_course_ method.

## 3. Materials and Methods

### 3.1. Simulation of canopy height data

For phenotyping experiments, one aims extracting (growth) parameter that are genotype-specific. In this work we used a simulation to demonstrate the suitability of dose-response models to extract temperature response parameters. The main advantage of using a simulation instead of real data is that one can report the expected precision and potential modeling artifacts. For real data experiments, judging on the heritability of extracted parameters would be the only choice, which does not allow distinguishing between poor performance caused by modeling artifacts and poor performance caused by lack of genetic effects.

As base for the simulation, we used the Wang-Engel function that has a similar shape as the Arrhenius function but is based on cardinal temperatures where one can rely on corresponding literature for temperature ranges (see Section Background). This approach allows examining whether simpler functions such as the asymptotic or linear function allow extracting temperature response parameters as well, despite the fact that they simplify the underlying ‘biological’ function.

The canopy growth of wheat genotypes was simulated using a growth function *g_T_* that was based on the Wang-Engel dose-response model of Equation 2 parameterized with four genotype-specific dose-response crop model parameters *θ^C^* = (*T*_min_, *T*_opt_, *T*_max_, *r*_max_). To mimic the characteristic variation in start and stop of the elongation phase for a diverse genotype set, this model was extended with time points for the start and end of growth, parameterized with two genotype-specific timing traits *θ^T^* = (*t*PH_start_, *t*PH_stop_). Due to this genotype-specific start and end of the growth process, values for measurement time points at the beginning and end of the season were unbalanced, potentially leading to modeling artifacts. An environment inhomogeneity factor *u* and a measurement error *e* were added, leading to the model

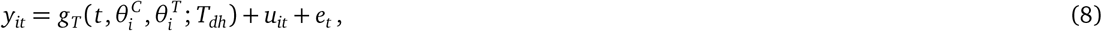

where *y_it_* is the measured canopy height (phenotype) for the *i*th genotype at campaign time points *t* in intervals of three days (*t* = 1, 4, 7,…, *T*), 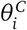 and 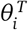 are genotype-specific crop growth model parameters respectively timing traits, *T_dh_* are hourly (*h*) temperature readings nested in measurement days (*d*), *u_it_* a time point and genotype-specific inhomogeneity error (i.e., simulating the influence of other covariates than temperature) and *e_t_* a time point specific measurement error (i.e., simulating random measurement errors). The growth function *g_T_* was specified as an implementation of Equation 7 based on the Wang-Engel function of Equation 2,

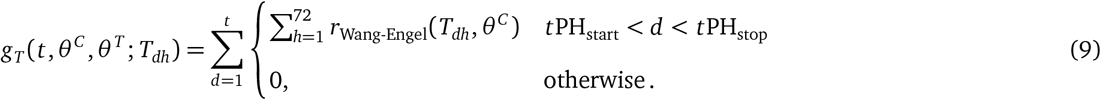

The genotype-specific traits 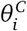 and 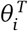 and the error *e_t_* were simulated using independent (genotypic) effects with variance *σ*^2^, and the field and environment imhomogeneities error *u_it_* according to a first-order autoregressive (AR1) model, all assuming normality. Specific *μ* and *σ*^2^ were chosen based on literature if available, and otherwise based on own field data (Roth, 2021). Corresponding sources and selected distributions for all simulation input parameter are summarized in Table 1. Two alternative simulations were performed: One with changing cardinal temperatures with time (indicated with ‘→’ in Table 1), and one with fixed cardinal temperatures (set to the mean of changing cardinal temperatures indicated in Table 1). Changing cardinal temperatures *T*_min_ and *T*_opt_ were implemented as linear interpolation between the start of the growth phase *t*PH_start_ with the corresponding lower cardinal temperate limit (relating to ‘terminal spikelet’ in Porter and Gawith (1999)), and the end of the growth phase *t*PH_stop_ with the corresponding upper cardinal temperature limit (relating to ‘anthesis’ in Porter and Gawith (1999)). Changing cardinal temperatures *T*_max_ were implemented as 5 °C ramp between the start of the growth phase *t*PH_start_ and the end of the growth phase *t*PH_stop_ from 30 to 35 °C (moving around the *T*_max_ of 31 °C reported for ‘anthesis’ in Porter and Gawith (1999), as precise values for ‘terminal spikelet’ are not provided).

**Table 1:**
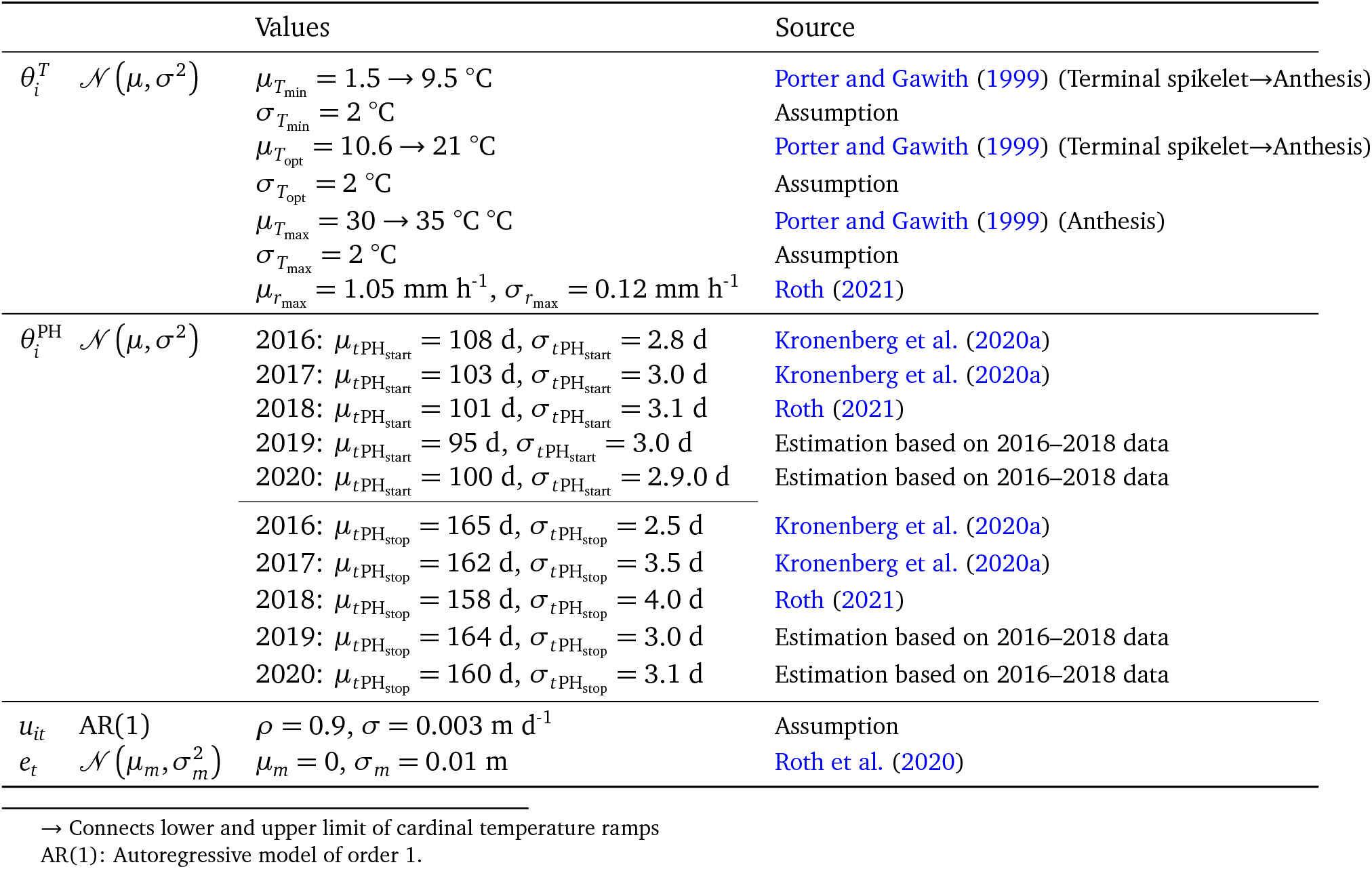
Model input parameters for the simulation.

|  | Values | Source |
| --- | --- | --- |
| $\theta_i^T \quad \mathcal{N}(\mu, \sigma^2)$ | $\mu_{T_{\min}} = 1.5 \rightarrow 9.5 \text{ }^\circ\text{C}$<br>$\sigma_{T_{\min}} = 2 \text{ }^\circ\text{C}$<br>$\mu_{T_{\text{opt}}} = 10.6 \rightarrow 21 \text{ }^\circ\text{C}$<br>$\sigma_{T_{\text{opt}}} = 2 \text{ }^\circ\text{C}$<br>$\mu_{T_{\max}} = 30 \rightarrow 35 \text{ }^\circ\text{C}$<br>$\sigma_{T_{\max}} = 2 \text{ }^\circ\text{C}$<br>$\mu_{r_{\max}} = 1.05 \text{ mm h}^{-1}, \sigma_{r_{\max}} = 0.12 \text{ mm h}^{-1}$ | Porter and Gawith (1999) (Terminal spikelet→Anthesis)<br>Assumption<br>Porter and Gawith (1999) (Terminal spikelet→Anthesis)<br>Assumption<br>Porter and Gawith (1999) (Anthesis)<br>Assumption<br>Roth (2021) |
| $\theta_i^{\text{PH}} \quad \mathcal{N}(\mu, \sigma^2)$ | 2016: $\mu_{t\text{PH}_{\text{start}}} = 108 \text{ d}, \sigma_{t\text{PH}_{\text{start}}} = 2.8 \text{ d}$<br>2017: $\mu_{t\text{PH}_{\text{start}}} = 103 \text{ d}, \sigma_{t\text{PH}_{\text{start}}} = 3.0 \text{ d}$<br>2018: $\mu_{t\text{PH}_{\text{start}}} = 101 \text{ d}, \sigma_{t\text{PH}_{\text{start}}} = 3.1 \text{ d}$<br>2019: $\mu_{t\text{PH}_{\text{start}}} = 95 \text{ d}, \sigma_{t\text{PH}_{\text{start}}} = 3.0 \text{ d}$<br>2020: $\mu_{t\text{PH}_{\text{start}}} = 100 \text{ d}, \sigma_{t\text{PH}_{\text{start}}} = 2.9.0 \text{ d}$<br>2016: $\mu_{t\text{PH}_{\text{stop}}} = 165 \text{ d}, \sigma_{t\text{PH}_{\text{stop}}} = 2.5 \text{ d}$<br>2017: $\mu_{t\text{PH}_{\text{stop}}} = 162 \text{ d}, \sigma_{t\text{PH}_{\text{stop}}} = 3.5 \text{ d}$<br>2018: $\mu_{t\text{PH}_{\text{stop}}} = 158 \text{ d}, \sigma_{t\text{PH}_{\text{stop}}} = 4.0 \text{ d}$<br>2019: $\mu_{t\text{PH}_{\text{stop}}} = 164 \text{ d}, \sigma_{t\text{PH}_{\text{stop}}} = 3.0 \text{ d}$<br>2020: $\mu_{t\text{PH}_{\text{stop}}} = 160 \text{ d}, \sigma_{t\text{PH}_{\text{stop}}} = 3.1 \text{ d}$ | Kronenberg et al. (2020a)<br>Kronenberg et al. (2020a)<br>Roth (2021)<br>Estimation based on 2016–2018 data<br>Estimation based on 2016–2018 data<br>Kronenberg et al. (2020a)<br>Kronenberg et al. (2020a)<br>Roth (2021)<br>Estimation based on 2016–2018 data<br>Estimation based on 2016–2018 data |
| $u_{it} \quad \text{AR}(1)$ | $\rho = 0.9, \sigma = 0.003 \text{ m d}^{-1}$ | Assumption |
| $e_t \quad \mathcal{N}(\mu_m, \sigma_m^2)$ | $\mu_m = 0, \sigma_m = 0.01 \text{ m}$ | Roth et al. (2020) |

→ Connects lower and upper limit of cardinal temperature ramps
AR(1): Autoregressive model of order 1.

Canopy growth was simulated for a measurement interval of three, seven and 14 days for a period between the 15th of March and 20th of July based on temperature courses of three years (Figure 2). Additionally, an irregular measurement frequency of two measurements per week with frequent shifts by one (*P* (1) = 1/3), two (*P*(2) = 2/9) or three (*P*(3) = 1/9) days was simulated. A total of 1,000 differing genotypes were assumed, resulting in 5,000 genotype time series. As temperature data source, real weather data of five consecutive years at the ETH research station of agricultural sciences in Lindau Eschikon, Switzerland (47.449 N, 8.682 E, 556 m a.s.l.) were used.

**Figure 2:**
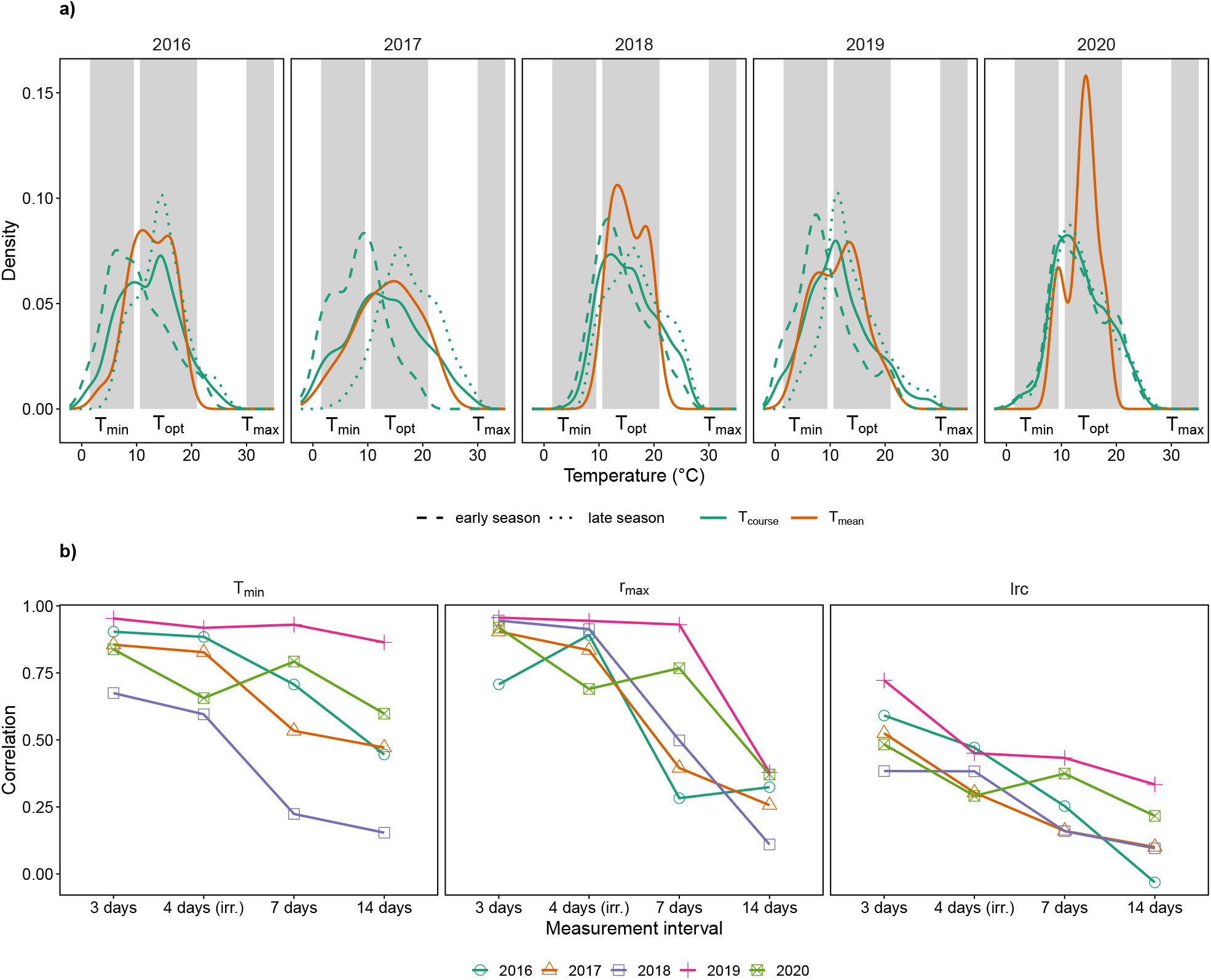
Measured temperatures in the stem elongation phase of winter wheat at the field phenotyping platform site of ETH Zurich (FIP) on an hourly scale (T_course_) and aggregated to a 3-day scale (T_mean_) (a) and Pearson’s correlation between simulation input and extracted parameter values in dependence on the measurement interval (b). Grey boxes in (a) indicate cardinal temperature ranges from terminal spikelet to anthesis according to Porter and Gawith (1999), early season corresponds to the first half of the stem elongation phase and late season to the second half. 4 days (irr.) in (b) corresponds to an intended measurement frequency of two measurements per week with frequent shifts by one, two or three days due to averse weather conditions, resulting in an irregular measurement frequency.

### 3.2. Extracting dose-response curves

To extract the Wang-Engel dose-response curve parameters *T*_min_, *T*_opt_ and *r*_max_, the linear and asymptotic growth response models (Equation 1 and 3) were fitted to canopy height data between *t*PH_start_ and *t*PH_stop_ for two covariate options, *T*_mean_ based on averaged temperatures between measurements (Equation 6), and *T*_course_ based on temperature courses with a period of one hour (Equation 7). In addition to the linear and asymptotic model, Equation 3 was enhanced with a slope parameter for *T*_min_ in dependence of time of measurement to test if changing cardinal temperatures can be integrated in the temperature response function,

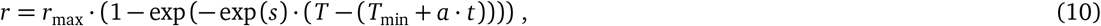

where *a* is the slope and *t* the day of measurement. Maximum-likelihood optimization was used to fit Equation 6 to a set of *t* data points, 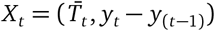. Fitting Equation 7 to data required modifying the definition of the data set so that 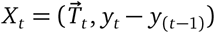, where 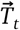 is a vector of covariate readings between trait measurement time points *t* – 1 and *t*.

As an independent random measurement errors was assumed, the parameter *σ* was fixed to the well-known measurement error of Structure-from-Motion based height estimations (Roth et al., 2020), herein the simulation input *σ_m_* (Table 1). Parameters were optimized using the method L-BFGS-B of the base R function *optim*.

To allow examining the effect of noise caused by inhomogeneities and measurement errors, the extraction was once performed on raw simulated data, once on data contaminated with the measurement error *e*, and once with both measurement and inhomogeneities errors *e* and *u*. Bias, variance, root-mean squared error (RMSE) and Pearson’s correlation were calculated after extracting the parameters *T*_min_, *lrc* and *r*_max_ for the asymptotic model and *a* (slope) and *T*_min_ for the linear model. Bias was defined as 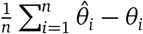, RMSE as 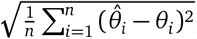, where *n* is the number of simulated time series, 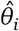 the extracted parameter value for the *i*th time series and *θ_i_* the simulation input parameter value.

### 3.3. Real field data case study

To give an outlook on the usefulness of combining high-resolution (hours) covariate data with low-resolution (days) HTFP data, the asymptotic growth response model and the novel *T*_course_ approach were applied to terrestrial laser scanning height data published in Kronenberg et al. (2020a). Three genotypes were selected (Nara, CH Claro, Suretta; Agroscope (Nyon, Switzerland) / DSP (Delley, Switzerland)). These genotypes were used as spatial checks in 2017 and therefore replicated 18 times, thus resulting in 54 plot based time series.

## 4. Results

Aggregating measured temperatures at the field phenotyping platform site of ETH Zurich (FIP) to hourly means (*T*_course_) and 3-day means (*T*_mean_) reveled severe differences in their distribution (Figure 2a): Temperatures close to the cardinal temperature *T*_min_ and between the cardinal temperatures *T*_opt_ and *T*_max_ were frequent for *T*_course_ in all years, but almost absent for *T*_mean_ in the years 2018 and 2020, and partly absent in 2016. While 2018 was a year with high temperatures throughout the stem elongation, 2020 was characterized by high temperatures in the early elongation phase. 2016 and 2017 were years with distinct temperature differences between the early and late season, while differences for 2019 were more moderate.

Simulating the stem elongation by combining the *T*_course_ measurements with a Wang-Engel dose-response function resulted in an average final height slightly higher than observed for elite varieties under field conditions by Kronenberg et al. (2020b) (Appendix, Figure 7), but realistic individual plant height time series with characteristic starts, stops, and lag phases (Figure 3). Fitting the asymptotic model (Equation 3) based on *T*_course_ (Equation 7) to the simulated data converged for 97% of all genotypes, fitting the model to real data of three genotypes with 18 replications for 91% of all plots. Based on visual inspections of simulated versus fitted curves (Figure 4a) the asymptotic model was able to model well the Wang-Engel curve increase in growth rate from *T*_min_ to *T*_opt_ while systematically underestimating *r*_max_ and slightly overestimating *T*_min_. Parameters extracted from real data were in the range of simulation input values with slightly lower *r*_max_ and higher *T*_min_ values (Figure 4b). Even tough for *T*_min_ the extracted values for the three genotypes were very close, significant differences between genotypes for all three parameters *T*_min_, *r*_max_ and *lrc* were found.

**Figure 3:**
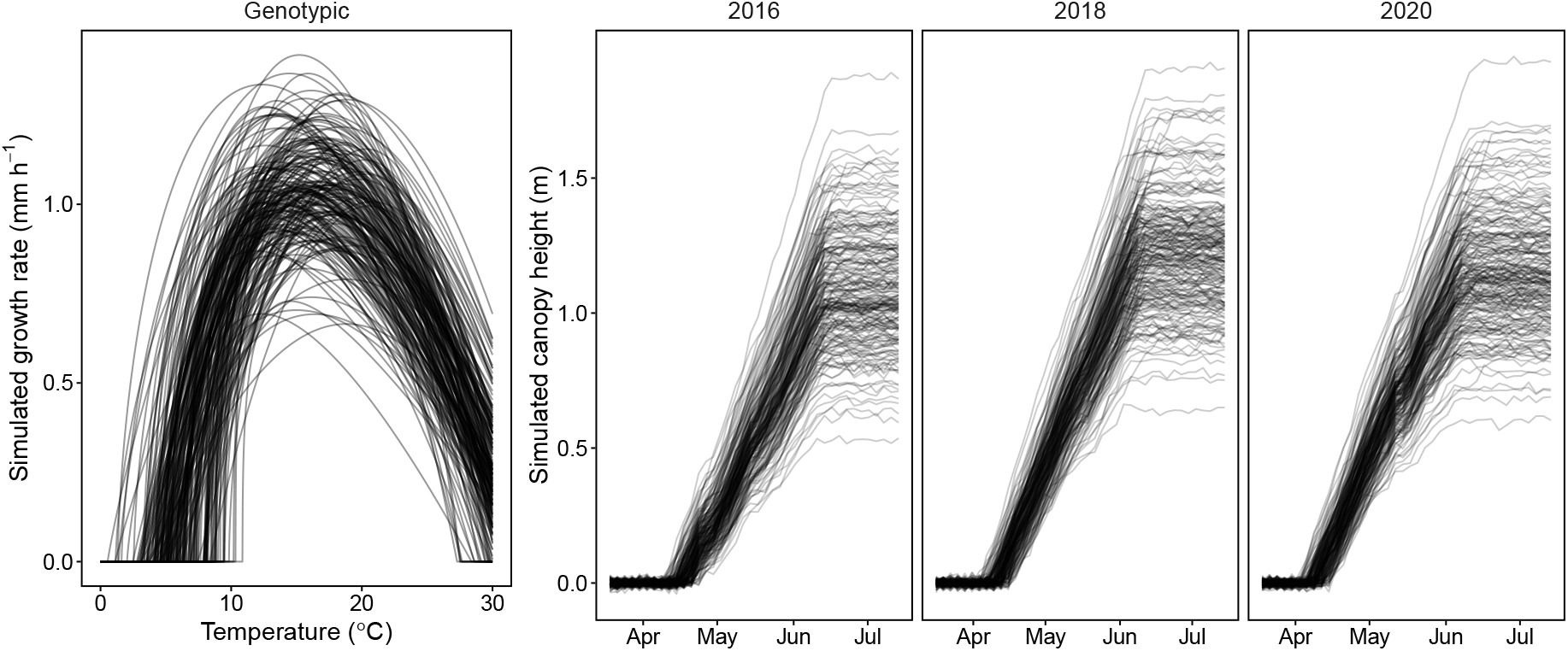
Simulated growth rates (a) and canopy heights (b) for 200 out of 1000 simulated genotypes and three years. Simulated growth rates are based on a Wang-Engel dose-response curve model.

**Figure 4:**
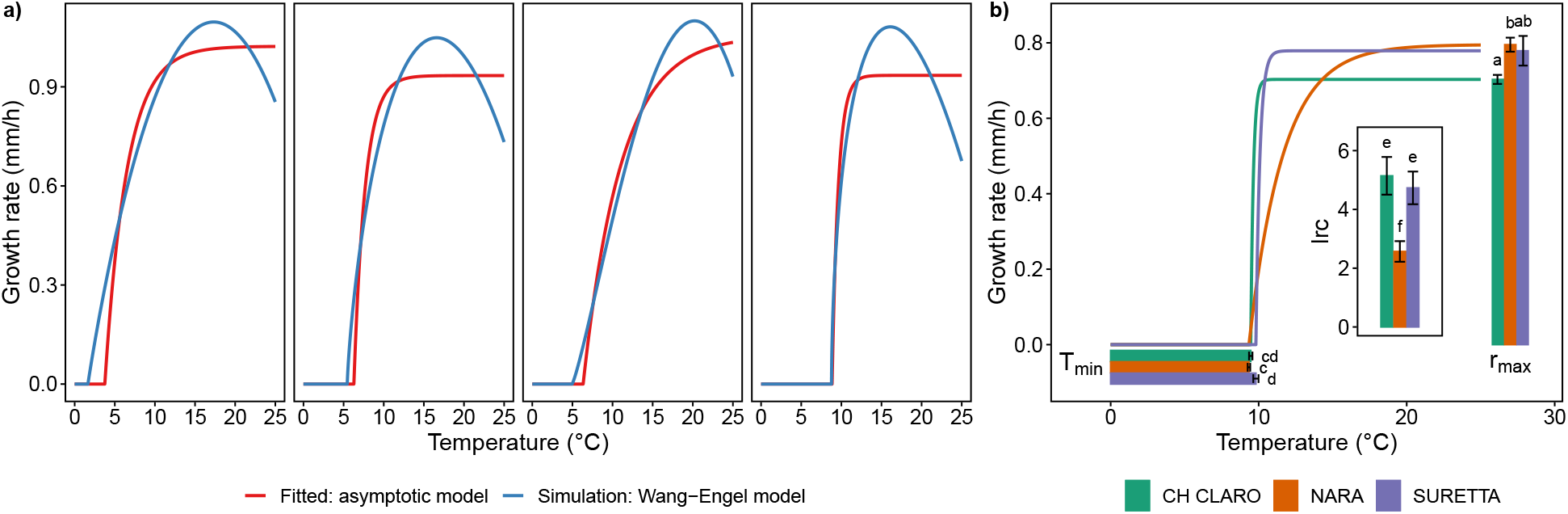
Simulated Wang-Engel dose-response curves and fitted asymptotic dose-response curves for four selected simulated genotypes (a), and real field data example with asymptotic curves fitted to TLS height data (terrestrial laser scanner) (Kronenberg et al., 2017) of three genotypes in 2017 (b). Lines and bars in (b) denote means of eighteen replications per genotype, error bars standard errors, letters significant differences (*α* = 0.05) based on one-way ANOVAs and Tukey’s Honest Significant Difference method.

Fitting the cardinal temperature shift enhanced model (Equation 10) to data converged for 90% of all genotypes. All other models converged for all genotypes. In a preliminary run, we also tried to fit the Wang-Engel model to simulated data, but failed to extract meaningful parameters. This was partly due to the complexity of defining valid non-overlapping ranges for the cardinal temperatures, but mainly because the model did not converge in most cases.

Based on the RMSE, the base temperature *T*_min_ were best estimated with the asymptotic dose-response model based on *T*_course_ covariates, while the maximum absolute growth rate *r*_max_ was best estimated with the asymptotic dose-response model based on *T*_mean_ covariates (Table 2). While the variance was drastically lower for both parameters if using *T*_course_, the bias slightly increased for *r*_max_ but decreased for *T*_min_. The RMSE, bias and variance of the linear model for the parameter *T*_min_ was much higher than for the asymptotic model.

**Table 2:**
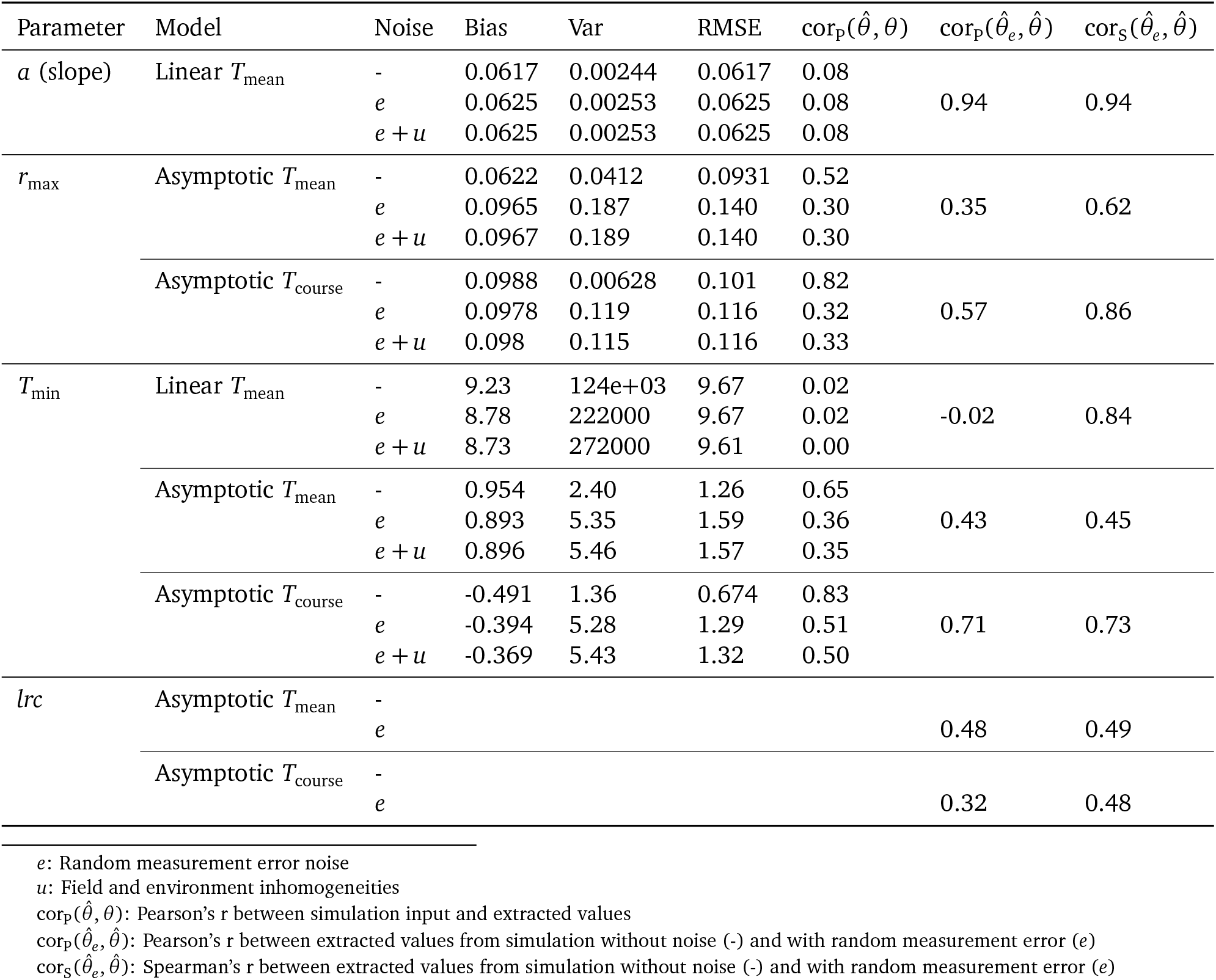
Bias, variance, root-mean squared error, and correlations for the linear and asymptotic model based on mean temperature and temperature courses for simulated data without noise, with noise due to measurement errors, and with noise due to field and environment inhomogeneities. Reported correlations are Pearson’s correlation between parameters extracted on simulated data and simulation inputs, and Pearson’s and Spearman’s correlation of the extracted parameters with and without noise due to measurement errors.

| Parameter | Model | Noise | Bias | Var | RMSE | $\text{cor}_p(\hat{\theta}, \theta)$ | $\text{cor}_p(\hat{\theta}_e, \hat{\theta})$ | $\text{cor}_s(\hat{\theta}_e, \hat{\theta})$ |
| --- | --- | --- | --- | --- | --- | --- | --- | --- |
| $a$ (slope) | Linear $T_{\text{mean}}$ | - | 0.0617 | 0.00244 | 0.0617 | 0.08 | | |
| | | $e$ | 0.0625 | 0.00253 | 0.0625 | 0.08 | 0.94 | 0.94 |
| | | $e + u$ | 0.0625 | 0.00253 | 0.0625 | 0.08 | | |
| $r_{\text{max}}$ | Asymptotic $T_{\text{mean}}$ | - | 0.0622 | 0.0412 | 0.0931 | 0.52 | | |
| | | $e$ | 0.0965 | 0.187 | 0.140 | 0.30 | 0.35 | 0.62 |
| | | $e + u$ | 0.0967 | 0.189 | 0.140 | 0.30 | | |
| | Asymptotic $T_{\text{course}}$ | - | 0.0988 | 0.00628 | 0.101 | 0.82 | | |
| | | $e$ | 0.0978 | 0.119 | 0.116 | 0.32 | 0.57 | 0.86 |
| | | $e + u$ | 0.098 | 0.115 | 0.116 | 0.33 | | |
| $T_{\text{min}}$ | Linear $T_{\text{mean}}$ | - | 9.23 | 124e+03 | 9.67 | 0.02 | | |
| | | $e$ | 8.78 | 222000 | 9.67 | 0.02 | -0.02 | 0.84 |
| | | $e + u$ | 8.73 | 272000 | 9.61 | 0.00 | | |
| | Asymptotic $T_{\text{mean}}$ | - | 0.954 | 2.40 | 1.26 | 0.65 | | |
| | | $e$ | 0.893 | 5.35 | 1.59 | 0.36 | 0.43 | 0.45 |
| | | $e + u$ | 0.896 | 5.46 | 1.57 | 0.35 | | |
| | Asymptotic $T_{\text{course}}$ | - | -0.491 | 1.36 | 0.674 | 0.83 | | |
| | | $e$ | -0.394 | 5.28 | 1.29 | 0.51 | 0.71 | 0.73 |
| | | $e + u$ | -0.369 | 5.43 | 1.32 | 0.50 | | |
| $lrc$ | Asymptotic $T_{\text{mean}}$ | - | | | | | | |
| | | $e$ | | | | | 0.48 | 0.49 |
| | Asymptotic $T_{\text{course}}$ | - | | | | | | |
| | | $e$ | | | | | 0.32 | 0.48 |

$e$ : Random measurement error noise
$\text{cor}_p(\hat{\theta}, \theta)$ : Pearson's $r$ between simulation input and extracted values
$\text{cor}_p(\hat{\theta}_e, \hat{\theta})$ : Pearson's $r$ between extracted values from simulation without noise (-) and with random measurement error ( $e$ )
$\text{cor}_s(\hat{\theta}_e, \hat{\theta})$ : Spearman's $r$ between extracted values from simulation without noise (-) and with random measurement error ( $e$ )

Introducing a random measurement error *e* had little effect on the correlation for the slope estimated by the linear model; extracted values correlated almost 1:1 for the simulation with and without such noise (Table 2). In contrast, the Pearson’s r for *T*_min_ was close to zero, but the Spearman’s rank correlation revealed nevertheless a high robustness of the extracted ranking to noise. The asymptotic model was more susceptible to noise than the linear model, indicated by increasing RMSE, bias and variance for *T*_min_ and *r*_max_. Using temperature courses instead of mean temperatures was of benefit for the parameters *r*_max_ and *T*_min_.

Correlating simulated versus predicted parameter values revealed further differences between the methods (Table 2, Figure 5): While the asymptotic model based on *T*_course_ was able to extract *r*_max_ and *T*_min_ with very strong correlations to the input values, the same model based on *T*_mean_ yielded only moderate correlations for *r*_max_ and *T*_min_.

**Figure 5:**
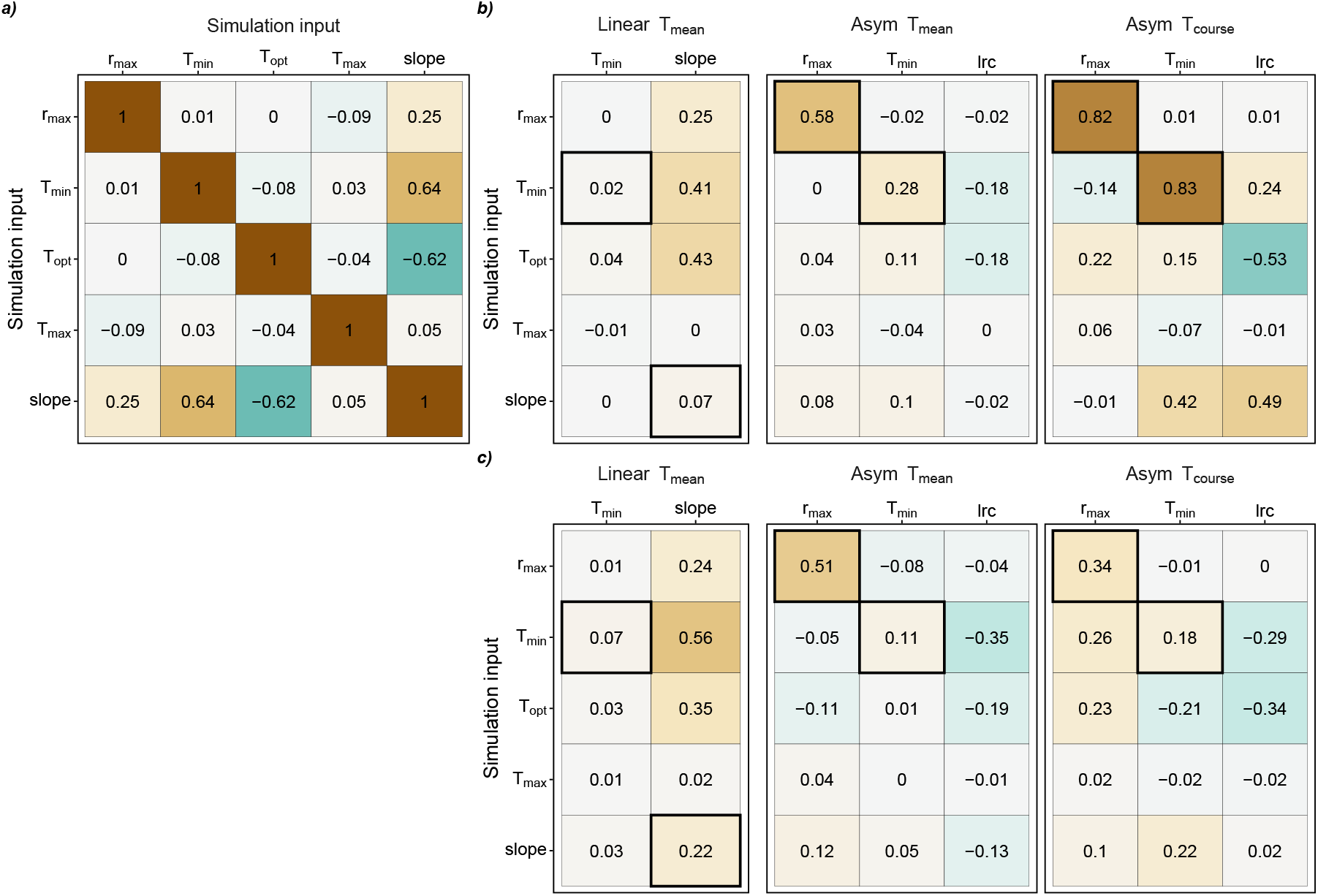
Pearson’s correlations of simulated data versus extracted temperature response parameters. Provided are results for the simulation without noise (a) versus extracted temperature response parameters with fixed cardinal temperatures (b) and changing cardinal temperatures with time (c) for the linear model based on mean temperature (Linear *T*_mean_) and the asymptotic growth model based on mean temperature (Asym *T*_mean_) and temperature courses (Asym *T*_course_). Note that the slope is derived from the relation slope = *r*_max_/*T*_opt_–*T*_min_ and therefore not an independent input parameter of the simulation. Black bold boxes indicate correlations between predicted and true values for identical parameters.

**Figure 6:**
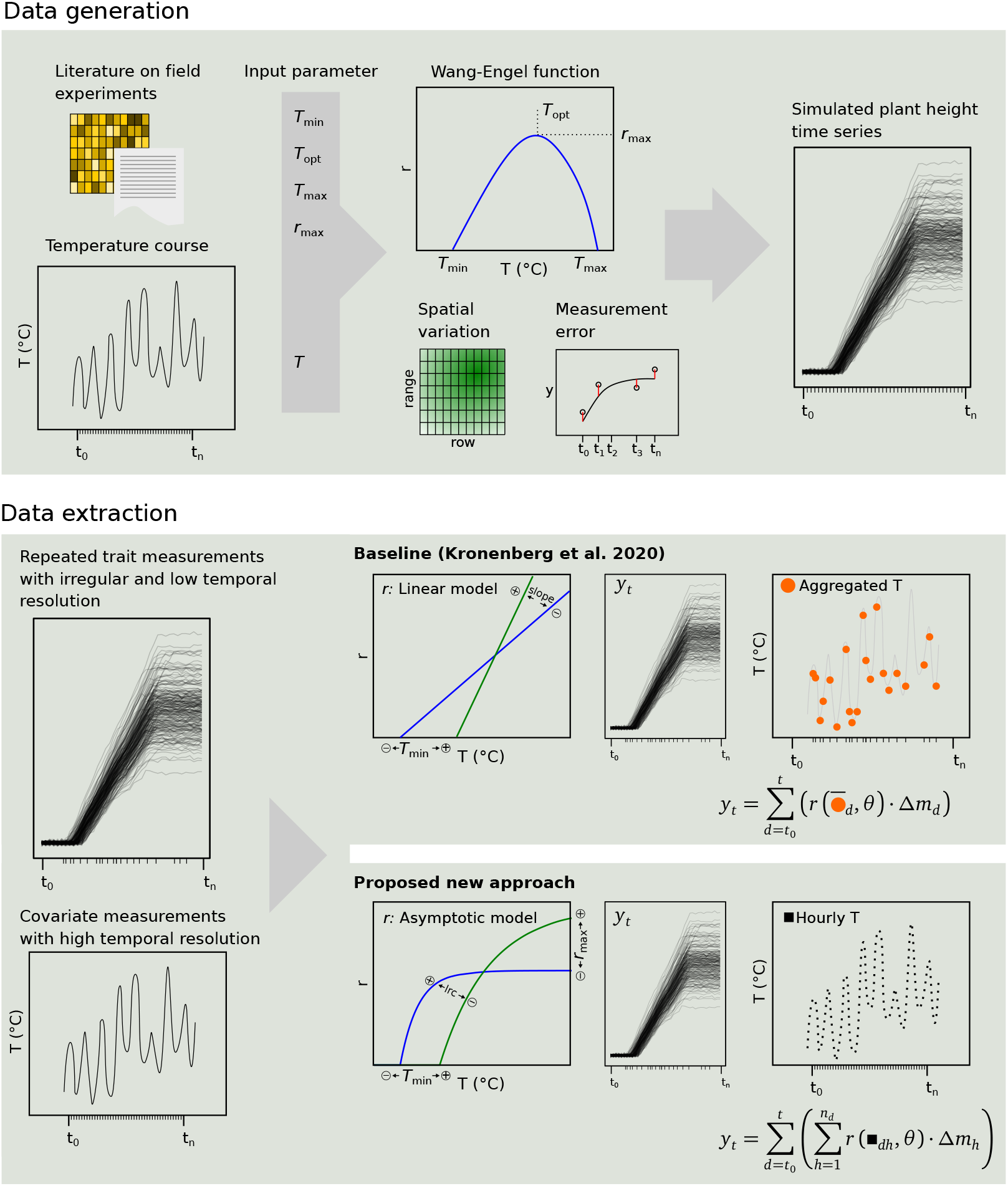
Schematic representation of the workflow. Top: Data generation using a simulation based on a Wang-Engel function parametrised with real temperature courses and cardinal temperatures from literature. Bottom: Data extraction using a linear model and aggregated temperature values (Kronenberg et al., 2020a) and the herein proposed new approach using an asymptotic model directly applied to covariate courses with high temporal resolution.

**Figure 7:**
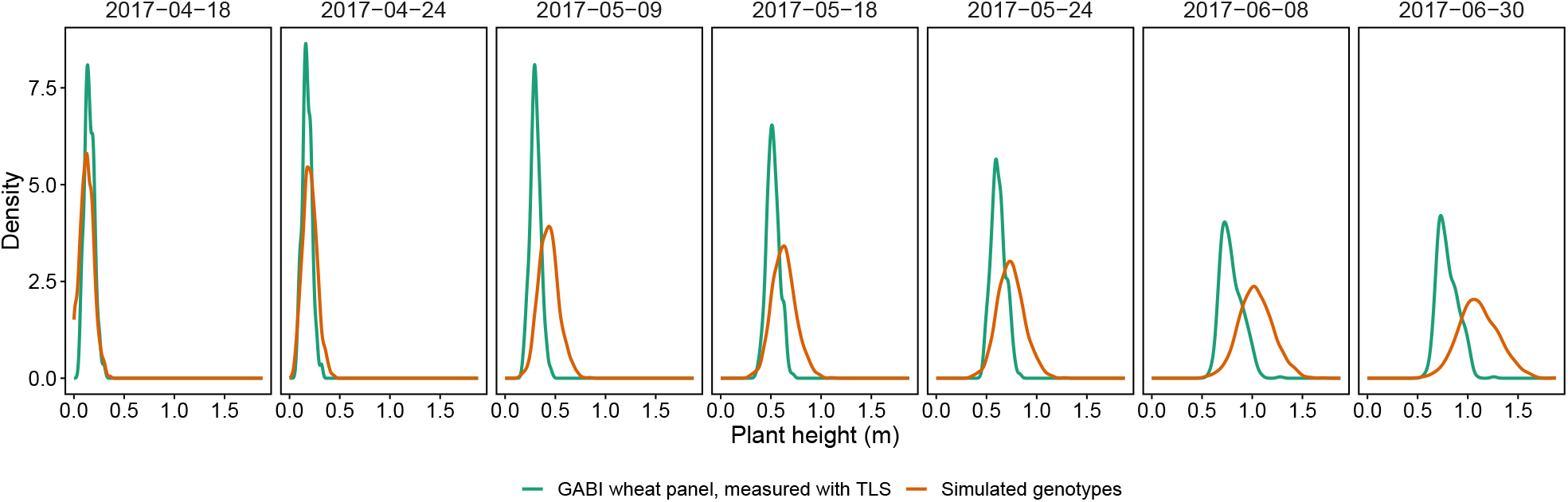
Simulated and measured plant heights for 2017. True measurement values were captured using a terrestrial laser scanner (Kronenberg et al., 2017) on a subset of the GABI wheat planel (350 genotypes). Simulated measurements values are based on 1000 assumed genotypes.

The parameter *r*_max_ extracted by the asymptotic model based on *T*_course_ was not only correlated with the simulated *r*_max_, but also weakly related to *T*_min_ and *T*_opt_ (Figure 5). Later correlations are at odds with the simulation input where *T*_min_, *T*_opt_ and *r*_max_ were uncorrelated, and is therefore an indicator of a modeling artifact. The asymptotic model additionally allowed estimating *T*_opt_ using *lrc* as proxy with a moderate correlation of *lrc* to *T*_opt_ for *T*_course_, while for *T*_mean_ the correlation was weak.

For the linear model, both parameter estimates of slope and *T*_min_ were uncorrelated with the corresponding simulated input parameter (Figure 5b). Instead, the slope was related to *r*_max_, *T*_min_ and *T*_opt_, while *T*_min_ based on the estimated intercept was not correlated to any input parameter. Both the linear and asymptotic model were not affected by supra-optimal temperatures: the input parameter *T*_max_ that defined growth in the supra-optimal range above *T*_opt_ was uncorrelated with any extracted parameter.

Adding noise to the simulated growth time series led to increased variances for the parameter estimates of *r*_max_ and *T*_min_ based on the asymptotic model (Table 2). While the RMSE increased by a factor of two for *T*_min_ and the correlation to input values dropped towards 0.5, it remained low for *r*_max_ while the correlation with input parameter dropped towards 0.3. The bias for *r*_max_ remained unchanged, while the bias for *T*_min_ decreased. Adding additional noise caused by highly auto-correlated spatial inhomogeneities did not further decrease the RMSE or increase the variance and bias.

Simulating changing cardinal temperatures for all genotypes alike decreased correlations of extracted parameters for all models (Figure 5c). The ranking for *T*_mean_ and *T*_course_ for the asymptotic model switched: While for both models the ability to extract *T*_min_ was drastically reduced, *T*_mean_ remained its ability to extract *r*_max_ but *T*_course_ lost it. The curvature parameter *lrc* was now moderately correlated with *T*_min_ and weakly with *T*_opt_ for *T*_mean_, therefore reducing its suitability as proxy trait for *T*_opt_. For *T*_course_ the correlation of *lrc* with *T*_opt_ remained moderate, therefore retaining its suitability as proxy trait for *T*_opt_. The slope extracted by the linear model was moderately related to *T*_min_ and weakly to *r*_max_ and *T*_opt_. Again, *T*_max_ was uncorrelated with any extracted parameter.

Integrating changing cardinal temperatures in the parameter extraction model turned out to be difficult (Appendix, Table 3): For *r*_max_ no improvement in parameter correlation and only slight improvements in RMSE, bias and variances were found. For *T*_min_ the correlation with input values slightly increased but remained weak, and RMSE, bias and variances slightly improved.

**Table 3:** Bias, variance, root-mean squared error, and Pearson’s correlation 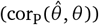 for values extracted from simulated data with noise and changing cardinal temperatures with the asymptotic model based on temperature courses (*T*_course_) and with an asymptotic model that additionally models the slope of changing cardinal temperatures (*T*_course&ramp_).

| Parameter | Model | Noise | Bias | Var | RMSE | $\text{cor}_p(\hat{\theta}, \theta)$ |
| --- | --- | --- | --- | --- | --- | --- |
| $r_{\text{max}}$ | Asymptotic $T_{\text{course}}$ | $u + e + \text{ramp}$ | 0.144 | 0.179 | 0.176 | 0.28 |
| | Asymptotic $T_{\text{course\&ramp}}$ | | 0.134 | 0.207 | 0.168 | 0.28 |
| $T_{\text{min}}$ | Asymptotic $T_{\text{course}}$ | $u + e + \text{ramp}$ | 1.91 | 11.2 | 2.21 | 0.16 |
| | Asymptotic $T_{\text{course\&ramp}}$ | | 1.86 | 10.6 | 2.20 | 0.23 |

The importance of the measurement interval varied with year and parameter (Figure 2b). Overall, 2019 was best suited to extract all parameters, independent of the measurement interval. For *T*_min_, in 2019 the measurement interval effect was almost negligible. For all other years, differences between three days intervals and four days irregular intervals were small for *T*_min_, while larger measurement intervals (7 and 14 days) severely reduced the correlations with input parameters. The same pattern, but even stronger reductions between four and seven days intervals, were visible for *r*_max_, except for 2019 where the correlation dropped only for a 14 days interval. For *lrc*, all reductions including the one from three days to four days reduced the correlations, and differences between the years were small.

## 5. Discussion

Although the growth of simulated canopies was based on a Wang-Engel temperature dose-response curve, using a corresponding model to extract parameters turned out to be not suitable, mainly because of failed convergence. Presumably, the low frequency of measurement values for supra optimal temperatures (≳ 30 °C, Figure 2) prevented estimating the required parameter *T*_max_. As suspected, the asymptotic model performed better with percentages of converged time series close to 100%. Using temperature courses instead of aggregated mean values allowed estimating independent parameters with higher correlations to input parameters, to the cost of a slightly decreased convergence rate. Especially the parameter *T*_min_ profited from taking temperature courses into account (Table 2), which indicates that in the early phase where stem elongation starts the distribution of temperatures and therefore temperature courses are essential to describe temperature responses (≲ 5 °C, Figure 2a). This thesis is also supported by the finding that years with a rather cold spring (2016, 2017, 2019) were better suited to extract *T*_min_ (Figure 2). Consequently, the use of the *T*_course_ method can provide dose-response curve models with valuable information about the occurrence of temperature values at the extremes, while the *T*_mean_ method leads to a shrinking of these values towards the mean.

Additionally to *T*_min_, the asymptotic model will extract the maximum absolute growth rate *r*_max_ and the parameter *lrc* which is related to steepness of the curvature of the asymptotic model. Accordingly, the parameter relates to the optimum temperature *T*_opt_. Thus, the model can fully describe temperature responses between the lower base temperature *T*_min_ and optimum temperature *T*_opt_. A limitation of the proposed asymptotic approach may be the high optimum temperatures: Although the extracted base temperature *T*_min_ and maximum growth rate *r*_max_ were strongly correlated with input parameters, they were also correlated to the optimum temperature *T*_opt_. As the parameters were simulated as uncorrelated, this indicates a modeling bias. Consequently, high optimum temperatures may lead to overestimation of *T*_min_ and *r*_max_

Noise and environment inhomogeneities reduced the correlation to input parameters drastically, independent on whether RMSE stayed stable or not (Table 2). It seems reasonable to assume that this reduction in correlation will lead to reduced estimated heritabilities of measured traits in HTFP. The simulation was not including a randomized field experiment set-up, but only independent genotype time series. Nevertheless, we assume based on our previous study on two other intermediate traits (Roth et al., 2021b) that also for dose-response curve traits it is essential to follow the three principles of experimental design, i.e., replication, randomization and blocking (Kempton, 1984). Such an experimental design will allow processing the extracted plot-based parameters using, e.g., linear mixed models to compensate for measurement errors and environment inhomogeneities and therefore ensure the overall heritability.

Simulating changing cardinal temperatures drastically reduced the ability of the asymptotic model to extract mean cardinal temperatures (Figure 5c). We suspect that this effect was overamplified by the simulation and that supposed cardinal temperature ramps in Porter and Gawith (1999) may be too extreme for wheat elite cultivar sets. In such an extreme ramp case, the asymptotic model would yield less heritable parameters than in the case of fixed cardinal temperatures, but would not introduce additional biases. Including an additional parameter to allow a shift of *T*_min_ with time only slightly improved the estimations (Appendix, Figure 3). In addition to a reduced convergence rate due to the increased number of parameters, introducing such a parameter also implies assuming the form of an underlying ‘biological’ function, which may further increases uncertainties as the exact form of the function is (yet) unknown. In addition to changes in cardinal temperatures, one may assume that *r*_max_ changes with time as well, but we are not aware of publications that would allow us to incorporate such changes in the simulation and extraction model.

A ‘close-to-optimum’ year 2019 was found in this study that was best suited to extract temperature dose-response parameters. The year was characterized by mild temperatures and clearly distinct temperature distributions for the early and late season of stem elongation. The early season covered well temperatures around *T*_min_, while late season temperatures ranged primarily around *T*_opt_ but also covered temperatures close to *T*_max_ (Figure 2a). In less optimal years, the chosen measurement interval plays an essential role (Figure 2b). Considering the fact that a field crop researcher cannot anticipate the suitability of a coming season to extract response parameters, an interval of at maximum 4 days is advisable.

The literature-based cardinal temperatures together with the sampled temperature courses during stem elongation suggest that *T*_opt_ but not *T*_max_ is reached at numerous hours. Thus, the asymptotic model proved to be superior compared to the more complex Wang-Engel model including *T*_max_, but also superior compared to the simpler linear model suitable for temperature ranges above *T*_min_ but well-below *T*_opt_. The former did not converge while the latter introduced substantial bias with regard to the estimation of cardinal temperatures. The linear response to temperature (slope) was similarly related to both cardinal temperatures *T*_min_ and *T*_opt_ but also to the maximum growth rate *r*_max_, but unrelated to the simulated slope (Figure 5b). Thus, the apparent temperature response is mainly driven by cardinal temperatures. When fitting a linear model to a wide temperature range, with significant number of hours in temperature ranges beyond the optimum (Figure 2a), it will inevitably compensate for both the maximum growth rate at the optimum and the base temperature of growth. Using a linear model if cardinal temperatures are changing during the season is even more harmful: The relation of the slope may switch from *T*_opt_ towards *T*_min_ (Figure 5c). Depending on the examined genetic material, applying a linear model may therefore lead to completely different conclusions.

Kronenberg et al. (2020a) applied such linear functions to wheat canopy height data. The authors argued that based on average temperatures in the measurement intervals, the temperature optimum *T*_opt_ of 27 °C (Parent and Tardieu, 2012) may not be reached. Nevertheless, our results show that the lower *T*_opt_ of 20.3 °C (Porter and Gawith, 1999) is reached at many hours during stem elongation, indicating that the slope reported by Kronenberg et al. (2020a) may be partly affected by growth at optimum temperature and the maximum growth rate. In contrast, the asymptotic model was able to mimic the curvature of a Wang-Engel model for temperature ranges below the optimum temperature (Figure 4a) and extracted differing values for *r*_max_ and *lrc* for three genotypes measured in the field (Figure 4b).

Whether an asymptotic model would also be more appropriate to model early canopy development (Slafer and Rawson, 1995; Grieder et al., 2015; Nagelmüller et al., 2016) remains questionable. Instead, the issue may rather be to model zero growth values below *T*_min_ appropriately. In summary, while a wide range of functions is available to model temperature-response, it is critical to know which cardinal temperature are reached under field conditions during the growth phase of interest, and consequently to use a function that is sufficiently flexible in critical areas of the affected temperature range.

## 6. Conclusion

Adequate models to quantify the different aspects of growth response to temperature may greatly improve our understanding of crop adaptation to certain climatic scenarios. Asymptotic models are a valid choice, if temperature ranges cover *T*_min_ and *T*_opt_ and the occurrence of supra-optimal temperatures during the observed growth phase is negligible. In contrast, linear models should be avoided in such situations. Using temperature courses with high temporal resolution as covariate input instead of aggregated mean temperatures is strongly advisable. The presence of changing cardinal temperatures—related to advancing phenology—may reduce the ability to extract parameters with an asymptotic model but will not severely bias results.

HTFP measurement intervals of about four days are sufficient to reliably extract *T*_min_ and *r*_max_ in an average season. Some irregularity in the frequency due to measuring breaks related to unsuitable weather conditions does not greatly affect the precision. Yet, *lrc* may profit from a higher measuring frequency. Not every season is equally suitable for characterizing dose-response curve parameters from repeated measurements, but an unsuitable season can be partially compensated for by increasing the measuring frequency. Consequently, we propose the following rule of thumb: “For dose-response modeling, one shall measure at least twice a week, but rest on weekends”.

## Model and Data availability

Data and source code that support the findings of this study are openly available in the ETH gitlab repository at https://gitlab.ethz.ch/crop_phenotyping/htfp_data_processing and archived in the ETH research collection (http://doi.org/10.5905/ethz-1007-385).

## Authorship contribution statement

**Lukas Roth**: Conceptualization, Methodology, Software, Formal analysis, Visualization, Writing - Original Draft. **Hans-Peter Piepho**: Conceptualization, Methodology, Writing - Review & Editing. **Andreas Hund**: Conceptualization, Supervision, Project administration, Funding acquisition, Writing - Review & Editing.

## Acknowledgements

We like to acknowledge Lukas Kronenberg for feedback on an early version of the manuscript, and Norbert Kirchgessner for feedback on formula notations (both ETH Zurich).

LR received funding from Innosuisse (http://www.innosuisse.ch) in the framework for the project “Trait spotting” (grant number: KTI P-Nr 27059.2 PFLS-LS). HPP was supported by DFG grant PI 377/24-1.

## Appendix

